# Why are amyloid-β plaques detected by X-ray phase-contrast imaging? Role of metals revealed through combined synchrotron infrared and X-ray fluorescence microscopies

**DOI:** 10.1101/2022.09.27.509706

**Authors:** Matthieu Chourrout, Christophe Sandt, Timm Weitkamp, Tanja Dučić, David Meyronet, Thierry Baron, Jan Klohs, Nicolas Rama, Hervé Boutin, Shifali Singh, Cécile Olivier, Marlène Wiart, Emmanuel Brun, Sylvain Bohic, Fabien Chauveau

## Abstract

Amyloid-β (Aβ) plaques from Alzheimer’s Disease (AD) can be visualized *ex vivo* in label-free brain samples using synchrotron X-ray phase-contrast tomography (XPCT). However, for XPCT to be useful as a screening method for amyloid pathology, it is essential to understand which factors drive the detection of Aβ plaques. The current study was designed to test the hypothesis that Aβ-related contrast in XPCT could be caused by the Aβ fibrils and/or by metals trapped in the plaques. This study probed the fibrillar and elemental compositions of Aβ plaques in brain samples from different types of AD patients and AD models to establish a relationship between XPCT contrast and Aβ plaque characteristics. XPCT, micro-Fourier-Transform Infrared spectroscopy and micro-X-Ray Fluorescence spectroscopy were conducted on human samples (genetic and sporadic cases) and on four transgenic rodent strains (mouse: APPPS1, ArcAβ, J20; rat: TgF344). Aβ plaques from the genetic AD patient were visible using XPCT, and had higher β–sheet content and higher metal levels than the sporadic AD patient, which remained undetected by XPCT. Aβ plaques in J20 mice and TgF344 rats appeared hyperintense on XPCT images, while they were hypointense with an hyperintense core in the case of APPPS1 and ArcAβ mice. In all four transgenic strains, β-sheet content was similar, while metal levels were highly variable: J20 (zinc and iron) and TgF344 (copper) strains showed greater metal accumulation than APPPS1 and ArcAβ mice. Hence, a positive contrast formation of Aβ plaques in XPCT images appeared driven by biometal entrapment.

**Graphical Abstract:** 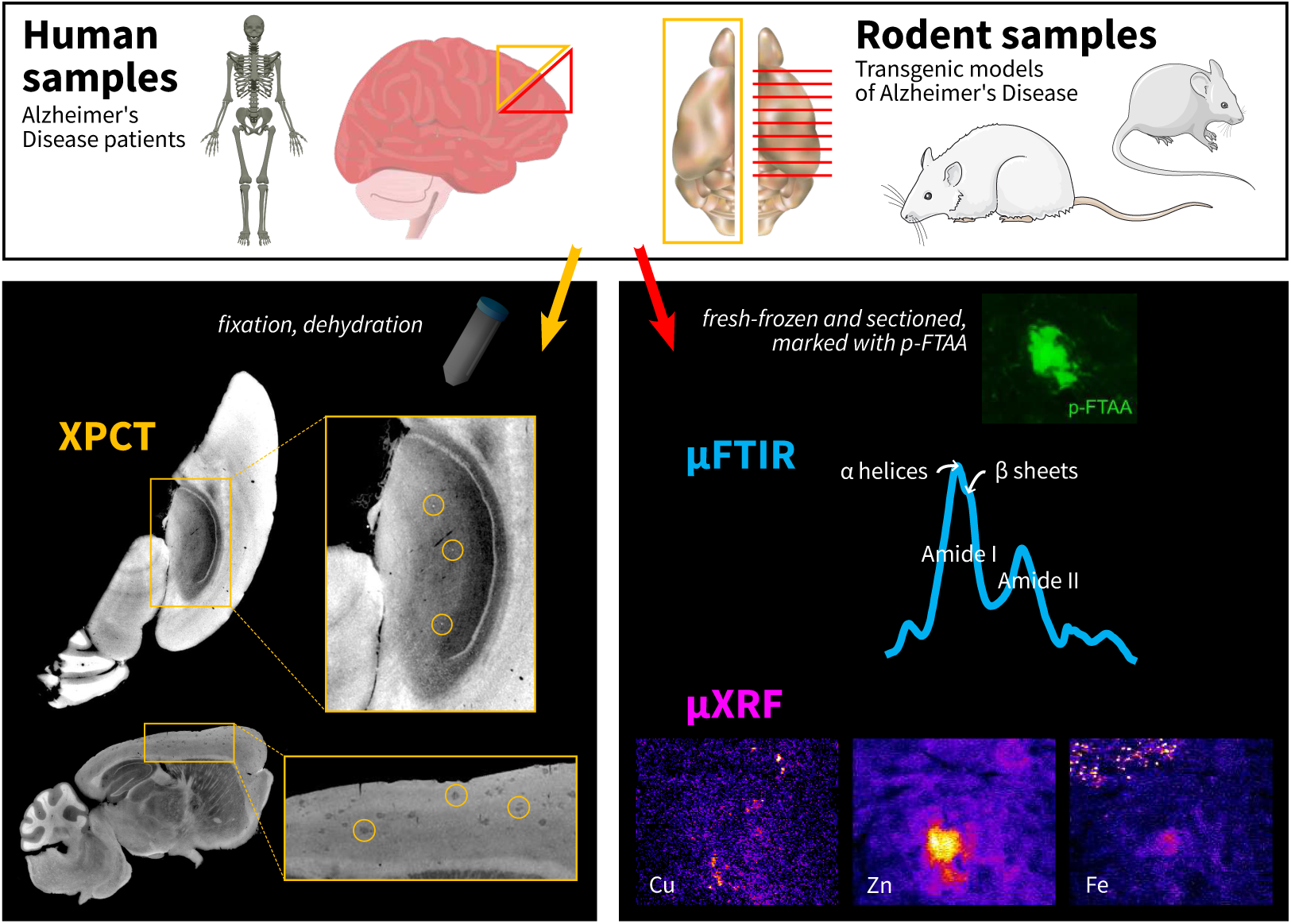

**Highlights:**

- Amyloid-β plaques in the different forms of Alzheimer’s Disease have various contrasts in X-ray phase-contrast tomography
- In transgenic rodents, a core-restricted, positive contrast is driven by the level of metal entrapment within plaques
- In humans, greater and more diffuse metal accumulation lead to a positive contrast in a genetic case of AD

## 1. Introduction

Amyloid-β (Aβ) plaques are an early hallmark of Alzheimer’s Disease (AD) in the human brain.Aβ plaques are formed by the aggregation of amyloid-β peptides with altered conformation into fibrils. Multiple techniques have arisen to assess the AD suspicion *in vivo*, especially Aβ positron emission tomography (PET) [1]. However, definite AD diagnosis still requires the *postmortem* examination of brain tissue and neuropathological scorings like the CERAD scoring [2], which qualitatively assesses the density and distribution within the brain, and Thal–Braak stages [3], which describe the progression of Aβ plaques within the brain areas.

X-ray phase-contrast tomography (XPCT) refers to a family of techniques which uses the phase shift rather than the attenuation of X-rays to probe the microstructure in soft biological tissues. X-ray 3D imaging of Aβ plaques has been described in transgenic rodent models that develop amyloidosis for the past decade, using propagation-based phase contrast (either with synchrotron radiation [4, 5, 6, 7] or with a laboratory source [8]) or using grating-based phase contrast (with synchrotron radiation [9, 10, 11, 4]). These techniques generate 3D images with an isotropic resolution ranging from 1 µm to 10 µm, thus providing a so-called “virtual histology” of the excised brain [12, 13, 14] with minimal preparation (fixation, dehydration or paraffin embedding). Imaging AD samples with XPCT is particularly interesting because plaques stand out in the images without the need to add any staining agents. Previous studies highlighted the great potential of XPCT for the quantitative analysis of amyloid-β plaques (overall burden [11, 4], spatial distribution [6], 3D morphometry [7]). These advances (which bring original information not available to standard immunohistochemistry), combined with increasingly faster acquisitions (less than 5 min per sample [7]), position XPCT as a potential screening tool for amyloid pathology. However, up to now, the substrate underlying contrast of Aβ plaques in XPCT images remains unexplored. In other words, it is not known why Aβ plaques are detected with XPCT.

Phase-contrast signal emerges from local changes of the X-ray refractive index within the plaques — compared to that of the surrounding tissue —, which makes them stand out. Changes of the refractive index could be caused by the characteristics of the fibrils structured as β-sheet stacks of amyloid-β peptides [15, 16, 17], which form dense and relatively insoluble aggregates. Or they could be linked to the entrapment of endogenous metals within the plaques, as iron, copper and zinc have been observed as co-localized with Aβ plaques, both in the human pathology [15, 18] and in rodent models [19, 20, 21]. We tested these two non-exclusive hypotheses to elucidate the origin of the phase-contrast signal of Aβ plaques. To this end, we performed an examination of the same excised brains from a variety of transgenic rodents (four transgenic strains) and AD patients (one sporadic case and one familial case) with three different and complementary synchrotron techniques, namely:

i. XPCT to visualize the plaques and measure the contrast with surrounding tissue,
ii. Fourier-transform infrared microspectroscopy (µFTIR) to quantify the proportion of β sheets in the plaques,
iii. X-ray fluorescence microspectroscopy (µXRF) to assess metal quantities in the plaques.

We here report new XPCT contrasts related to Aβ plaques: in rodents, some strains displayed a hypointense (dark) contrast while others had the already described hyperintense (bright) contrast. In humans, XPCT highlighted parenchymal Aβ plaques for the first time (in the genetic patient). Overall our results suggest that positive XPCT contrast is mainly driven by the level and spatial distribution of metal entrapment.

## 2. Material & Methods

### 2.1. Study design

Figure 1 summarizes study design, which intended to examine Aβ plaques from the same brains with XPCT, µFTIR and µXRF. Tissue preparation was not compatible between the three techniques, thus the samples had to be cut: one half was processed for XPCT, the other half was processed for µFTIR followed by µXRF. The list of samples, and all measurements performed on individual Aβ plaques (*n* = 9–23 per strain), is available as a supplementary file.

**Figure 1:**
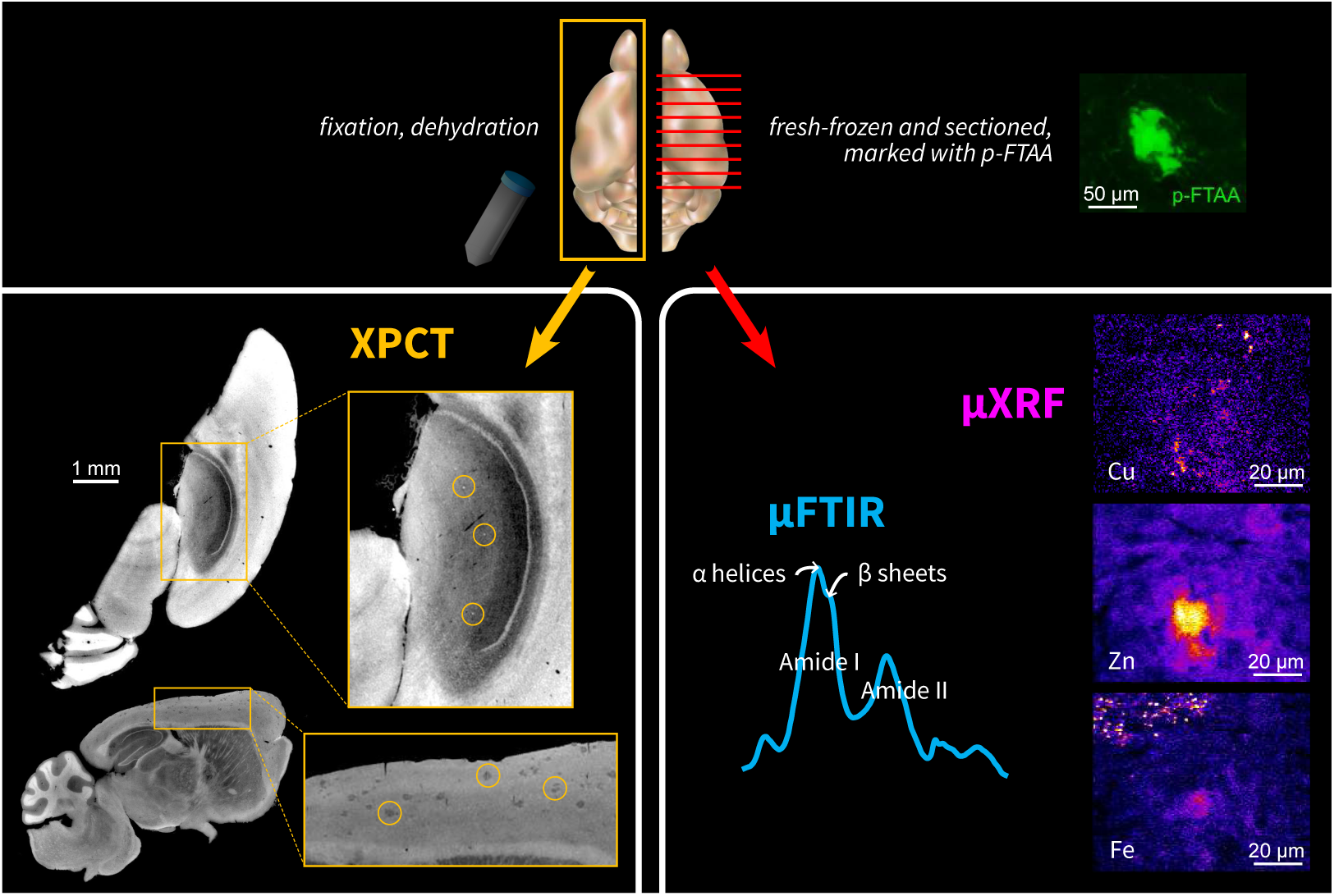
Experimental design, summarizing sample preparation and imaging techniques. A half of the sample was fixed and dehydrated then scanned in 3D with XPCT (left part). The other half was sectioned while frozen, fluorescently stained to locate Aβ plaques, and analyzed in 2D using µFTIR (for β-sheet load) and µXRF (for metal level).

### 2.2. Samples

#### 2.2.1. Human brain samples

Human brain samples (from parietal and frontal cortex) and associated data were obtained from Tissu-Tumorothèque Est Biobank (CRB-HCL Hospices Civils de Lyon) authorized by the French Ministry of Research (authorizations DC2008-72 & AC2015-2576):

- A 51-year-old male patient with genetic (APP duplication) AD (postmortem delay 36 h);
- An 82-year-old female patient with sporadic AD (post-mortem delay 11 h).

Both had pronounced Aβ pathology (Thal–Braak stage V, CERAD score 3) [22], and had been previously diagnosed as free from prion disease.

Two snap-frozen samples were collected. One sample was kept frozen for sectioning, while the other was fixed with formaldehyde 4 % (as described in the tissue preparation section below).

Paired formalin-fixed paraffin-embedded (FFPE) blocks were cut onto slides and stained with multiple agents:

- Anti-Aβ A4 antibodies for Aβ load;
- Anti-phosphorylated-tau antibodies for neurofibrillary tangles (NFTs);
- Luxol fast blue and periodic acid-Schiff (PAS) stains for myelin and glycogen-rich bodies.

#### 2.2.2. Transgenic rodent brain samples

Four transgenic rodent models that develop amyloidosis were studied:

- Mono-transgene line ArcAβ (*n* = 3 mice, 15 months old), with mutant APP [23] (https://www.alzforum.org/research-models/arcav)
- Double-transgene line APPPS1 (*n* = 3 mice, 12 months old), with mutant APP and PSEN1 [24] (https://www.alzforum.org/research-models/appps1)
- Mono-transgene line J20 (*n* = 2 mice, 13 months old), with mutant APP [25] (https://www.alzforum.org/research-models/j20-pdgf-appswind)
- Double-transgene line TgF344 (*n* = 3 rats, 20 months old), with mutant APP and PSEN1 [26] (https://www.alzforum.org/research-models/tgf344-ad)

One wild-type (WT) rat (Fischer 344) and one WT mouse (C57Bl/6), respectively 20 and 15 months old, were used as controls.

Rodents were sacrificed with intracardiac PBS perfusion to clean blood vessels. Brains were excised and kept at −80 °C.

Mouse brains were cut in half along the hemisphere (Figure 1). One half was then fixed and dehydrated (for XPCT). The other half was immediately put back at −80 °C (for µFTIR and µXRF).

Only one hemisphere was available per rat brain. The hemispheres were cut in the middle of the hippocampus. One half was then fixed and dehydrated (for XPCT). The other half was immediately put back at −80 °C (for µFTIR and µXRF).

#### 2.2.3. Fixed tissue and dehydration

The unfrozen samples were fixed for 24 to 48 hours with 4 % FA. Then they were dehydrated at RT in a series of 8 baths with a gradient of ethanol concentrations, diluted with PBS, for 5 minutes each: 25 %, 25 %, 50 %, 50 %, 75 %, 75 %, 96 %, 96 %. Finally, they were put into plastic tubes filled with 96 % ethanol and kept at 4 °C.

#### 2.2.4. Frozen tissue and preparation of SiRN membranes

The frozen samples were dissected into blocks in the hippocampus / upper cortex area, to ensure they would fit into an area of 4 × 4 mm^2^. Then they were sectioned using a Leica CM1850 cryostat at −20 °C with a thickness of 7 µm. PTFE-coated blades (DuraEdge™, TedPella, Inc., USA) were used to prevent metallic contamination. Brain sections were placed with a paintbrush on 200 nm-thick silicon-rich nitride (SiRN) membranes (Silson Ltd, UK) with an inner frame side of 4 millimeters. Sections were stained for 5 minutes with a pan-amyloid fluorescent agent, p-FTAA [27] (1 µmol L^−1^ in PBS with 0.01 % NaN_3_); membranes were then rinsed once with nanopure water. Fluorescence and bright-field transmission images were acquired with a ZEISS microscope (cf. Figure 2, A). Afterwards, membranes were kept in a desiccator with silica gel blue (2 mm–4 mm, Roth) until synchrotron experiments (1–4 weeks after).

**Figure 2:**
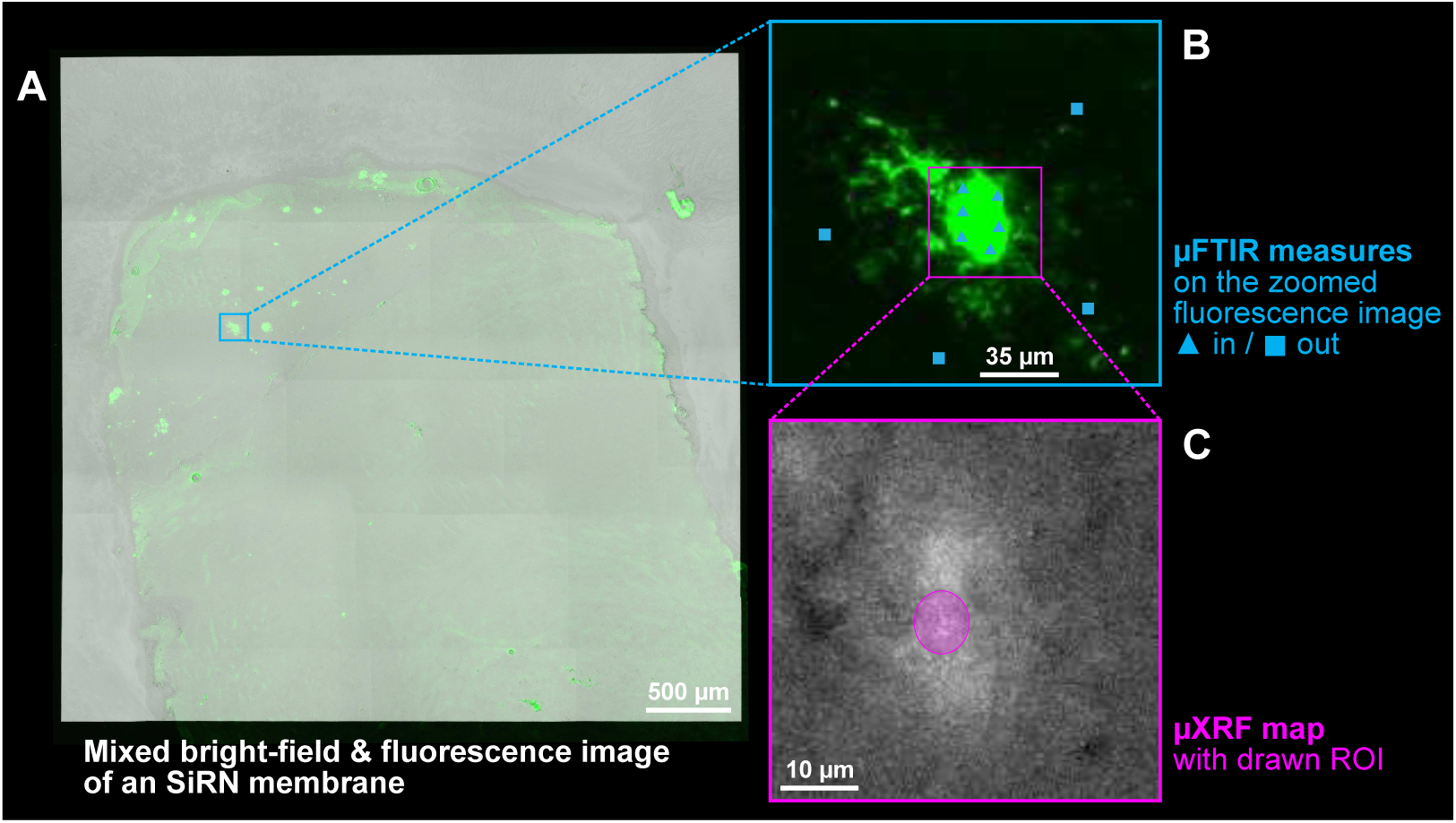
Silicon-nitride membrane overview and 2D imaging with µFTIR and µXRF. (A) Sectioned samples were placed on SiRN membranes, fluorescently stained and imaged with a microscope with 2 channels: bright-field and green fluorescence. (B) During the experiment at the SMIS beamline, the fluorescence was used to target Aβ plaques and multiple µFTIR spectra were acquired within individual plaques and in the surrounding tissue. (C) During the experiment at the NANOSCOPIUM beamline, µXRF hypermaps were acquired on the same Aβ plaques — here shown as the sum over the whole spectrum.

### 2.3. Synchrotron experiments

All experiments, unless explicitly stated, were conducted at the synchrotron radiation facility SOLEIL, within the scope of the proposal #20200258.

#### 2.3.1. X-ray phase-contrast tomography (XPCT)

Formaldehyde-fixed, ethanol-dehydrated samples were imaged with X-ray phase-contrast tomography (XPCT) at the ANATOMIX beamline [28] with a pink beam of mean energy 22 keV and a detector pixel size of 6.5 µm. The detector system consisted of a lutetium aluminum garnet (LuAG, Lu_3_Al_5_O_12_) single-crystal scintillator (600 µm thick) coupled to a CMOS camera (Hamamatsu Orca Flash 4.0 V2, 2048 × 2048 pixels) via a 1:1 lens system consisting of two Hasselblad photo objectives (*f* = 100 mm) in tandem geometry. Complete acquisition parameters are reported in Supplemental Table 1.

For reconstruction using PyHST2 [29], we set the Paganin length parameter to 552 µm (*δ/β* = 1525). We used a motion blur correction algorithm and an unsharp mask filter (*c_unsharp_* = 0.5, *σ_unsharp_* = 2 px).

Human samples were imaged at a higher resolution via a 1.45:1.45 lens system using a similar setup. The Paganin length was adapted to 216.65 µm (*δ/β* = 1400).

#### 2.3.2. XPCT image analysis

Ring artifacts were removed using an in-house tool implemented by the company NOVITOM (France; https://www.novitom.com/en/) [14] Samples were isolated from the background (air, plastic tube and ethanol) and rotated within the AMIRA 3D software (release 2021.1, Thermo Fisher Scientific, USA). Aβ plaques were segmented semi-automatically using a random forest algorithm from the IPSDK software (Reactiv’IP, France; https://www.reactivip.com/). The generated masks were screened for false positive segmentations and ∼ 20 plaques were selected, along with ROIs of the background. Signal intensity measurements — as a ratio of the mean intensities of the plaques and the mean intensities of the background ROIs — were retrieved with MorphoLibJ [30], an ImageJ plugin.

#### 2.3.3. Fourier-transform infra-red microspectroscopy (µFTIR)

The brain cryosections on the membranes (cf. Figure 2, panel B) were analyzed by synchrotron-radiation FTIR microspectroscopy with a Continuum XL microscope and a Thermo Nicolet 6700 bench at the SMIS beamline. The microscope was equipped with a fluorescence imaging setup and an halogen mercury lamp for excitation of the p-FTAA fluorescent probe that binds Aβ plaques.

Spectra on and around the spots of interest (respectively referred as “In” and “Out”) were recorded between 650 cm^−1^ and 4000 cm^−1^ at 4 cm^−1^ in 10 × 10 µm^2^ dual-aperture mode, using a Schwarzschild 32× (NA 0.65) objective and a matching condenser; zero-padding enhanced resolution to 0.5 cm^−1^. All parameters are shown in Supplemental Table 2. Spectra were pre-processed with the atmospheric suppression algorithm from OMNIC 9.2 (Thermo Fisher Scientific, USA) to remove water vapor contribution.

Besides, a few acquisitions were performed as evenly-spaced arrays (“maps”) with a lower co-addition to scan larger areas on a few samples, especially those devoid of Aβ from the WT animals.

#### 2.3.4. µFTIR spectral analysis

Spectra were processed using Quasar [31]. All the data were imported and processed with the same pipeline:

Cutting: The region from 650 cm^−1^ to 800 cm^−1^ was cut out because of high-frequency noise;

Quality assessment: Spectra were thoroughly screened for Mie scattering artifacts — indeed, Mie scattering is responsible for the shifting of the Amide I peak towards lower wave numbers and for its distortion, which leads to measurements that are easily mistaken for higher β-sheet content [32];

Absorbance thresholding: MCT detectors have a limited range for linearity between 0.2 and 1.4, so spectra with a peak-to-peak amplitude outside this range were discarded;

Baseline correction: A linear baseline correction was applied on the Amide I band, from 1415 cm^−1^ to 1770 cm^−1^;

Feature selection: Absorbance values closest to 1655 cm^−1^ (non-β sheets) and 1630 cm^−1^ (β sheets) [33] were used as features to compute a spectroscopic ratio for amyloid β-sheet content: 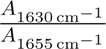

#### 2.3.5. X-ray fluorescence microscopy (µXRF)

The brain cryosections on the membranes (cf. Figure 2, panel C) were analyzed by x-ray fluorescence (µXRF) microscopy on the “CX3” custom bench at the NANOSCOPIUM beamline, using the “flyscan” mode with steps of 0.3 µm. Measures were performed on the same locations on the membranes as for µFTIR acquisitions (cf. § 2.3.3), through the method described in **??**. The inline optical bright-field microscope which is mounted on the beamline helped fine-tuning the position. All parameters are shown in Supplemental Table 3.

Three standard reference materials (SRMs) were also scanned: the SRM 1577c “bovine liver” and SRM 1832 (NIST, USA), along with the “RF” standard thin film (AXO DRESDEN GmbH, Germany).

#### 2.3.6. µXRF elemental analysis

Spectra were processed using PyMca [34]. First, the raw spectral maps were summed to locate the plaques more precisely within the scanned field of view — from 20 × 20 µm^2^ to 80 × 80 µm^2^. Regions of Interest (ROIs) were drawn at the location of each plaque.

Then, the measures on the three SRMs and on a blank SiRN membrane helped the calibration of the software — using the fundamental parameter method — and the batch-fitting tool computed the mass fraction maps for each element. Reported mass fractions in µg g^−1^ or “parts per million” (ppm) were obtained by averaging within the drawn ROI. One can easily retrieve an approximate surface concentration: 10 ppm ≈ 1 ng cm^−2^; because *e_dried tissue_* ≈ 1 µm and *ρ_dried tissue_* = 1.04 g cm^−3^ (the density of dried brain tissue).

### 2.4. Statistics

Results are displayed as box plots which use the Tukey standards, i.e. the whiskers end at *Q*1 − 1.5 × *IQR* and *Q*3 + 1.5 × *IQR* (*IQR* is the interquartile range); values outside of [*Q*1 − 1.5 × *IQR*; *Q*3 + 1.5 × *IQR*] are represented as points; means are represented as crosses and medians as a wide line. Reported fold increases are computed between the means.

The normality and log-normality D’Agostino-Pearson tests failed for the whole data set. Kruskal-Wallis tests with post-hoc Dunn’s multiple comparison tests were performed on XPCT intensities, µFTIR-derived data and µXRF-derived data in rodents. A separate Kruskal-Wallis test with post-hoc Dunn’s multiple comparison tests was performed for human µFTIR-derived data. For human µXRF-derived data, a non-parametric Mann-Whitney test was performed per element (iron, zinc and copper levels were considered independent).

## 3. Results

### 3.1. Genetic and sporadic human AD samples exhibit different Aβ contrasts in XPCT

On XPCT images (Figure 3), the multiple Aβ plaques from the genetic AD patient were clearly visible in grey matter, exhibiting a typical neuritic aspect (Figure 3, A1-B1, yellow arrow heads) and diameter around 50 µm. Anti-Aβ staining of slides derived from paired FFPE block confirmed the presence of numerous plaques with dense central core and less compact peripheral halo (Figure 3, C1, yellow arrows). The central core of neuritic plaques and surrounding dystrophic neurites containing neurofibrillary material were also stained using anti-p-tau antibodies (Figure 3, D1, yellow arrows). The presence of multiple Aβ plaques in this familial form of AD was especially striking when scrolling through the virtual sample (movie). In contrast, the numerous Aβ diffuse plaques detected in Aβ-stained slides from the sporadic AD patient could not be reliably visualized in XPCT (Figure 3, red arrow heads). In both genetic and sporadic AD cases, smaller, round-shaped and highly hyperintense structures were observed in sub-pial layer of frontal sulcus. These structures were presumably corporea amylacea, as far as similarly shaped inclusions positive to periodic acid-Schiff (PAS) staining were found in similar location in slides from FFPE paired blocks for both genetic and sporadic AD (Figure 3, blue arrows). Additional triangle shaped structures disseminated across cortex layers were also seen. They were enhanced after Maximum Intensity Projection (MIP). Their shape and distribution across the cortex corresponded to p-tau-stained tangles in FFPE-derived slides (Figure 3, green arrows). Other variably hyperdense structures were neuronal bodies and vessel walls.

**Figure 3:**
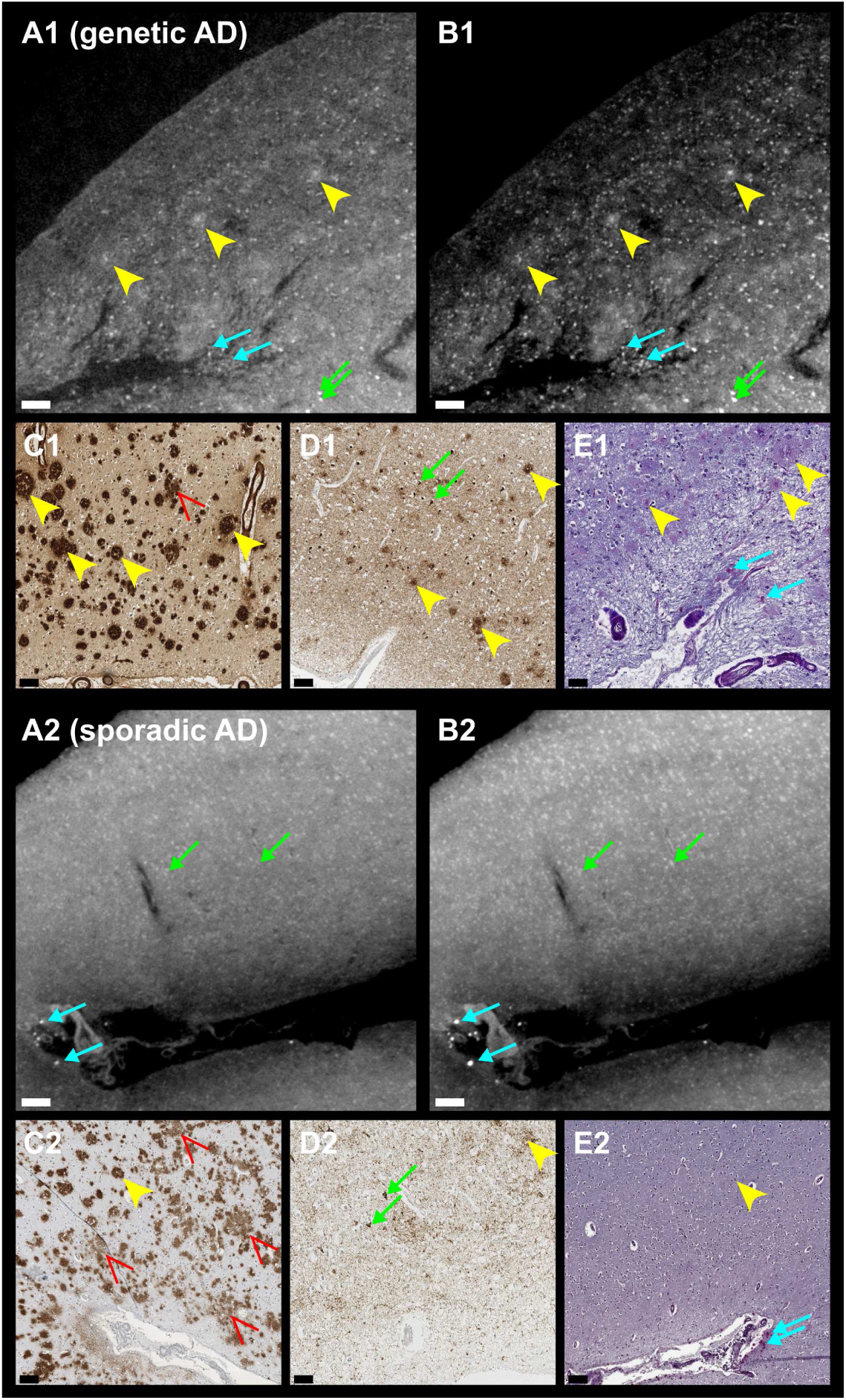
Representative XPCT images of brain tissue with Aβ plaques from a genetic (A1-E1) and a sporadic (A2-E2) cases of Alzheimer’s Disease. Insets A1&A2 are single slices while insets B1&B2 are maximum intensity projections (MIPs) of the same FOVs over 5 slices (16 µm). Virtual slices are compared to (contralateral) FFPE slices: anti-Aβ A4 immunostaining (C1&C2), anti-p-tau immunostaining (D1&D2) and LFB / PAS staining (E1&E2). Yellow arrow heads point at neuritic amyloid plaques while red arrow heads point at diffuse plaques. Other hyperdensities, smaller than Aβ plaques, were p-tau stained tangles (green arrows) and corporea amylacea (blue arrows). Neuron soma and blood vessel walls were slightly hyperdense. Scale bars equal 100 µm.

### 3.2. β-sheet content and metal entrapment are strikingly enhanced in the genetic case compared to the sporadic case

As expected, the β-sheet content in the fluorescence-labelled plaques (Figure 4, panel A) was higher than in the surrounding tissue for both patients (genetic AD: +50 %; sporadic AD: +4 %). Besides, the β-sheet content significatively differentiated Aβ plaques between the genetic and sporadic AD patients (genetic AD: +55 %; *p* = 0.0291).

**Figure 4:**
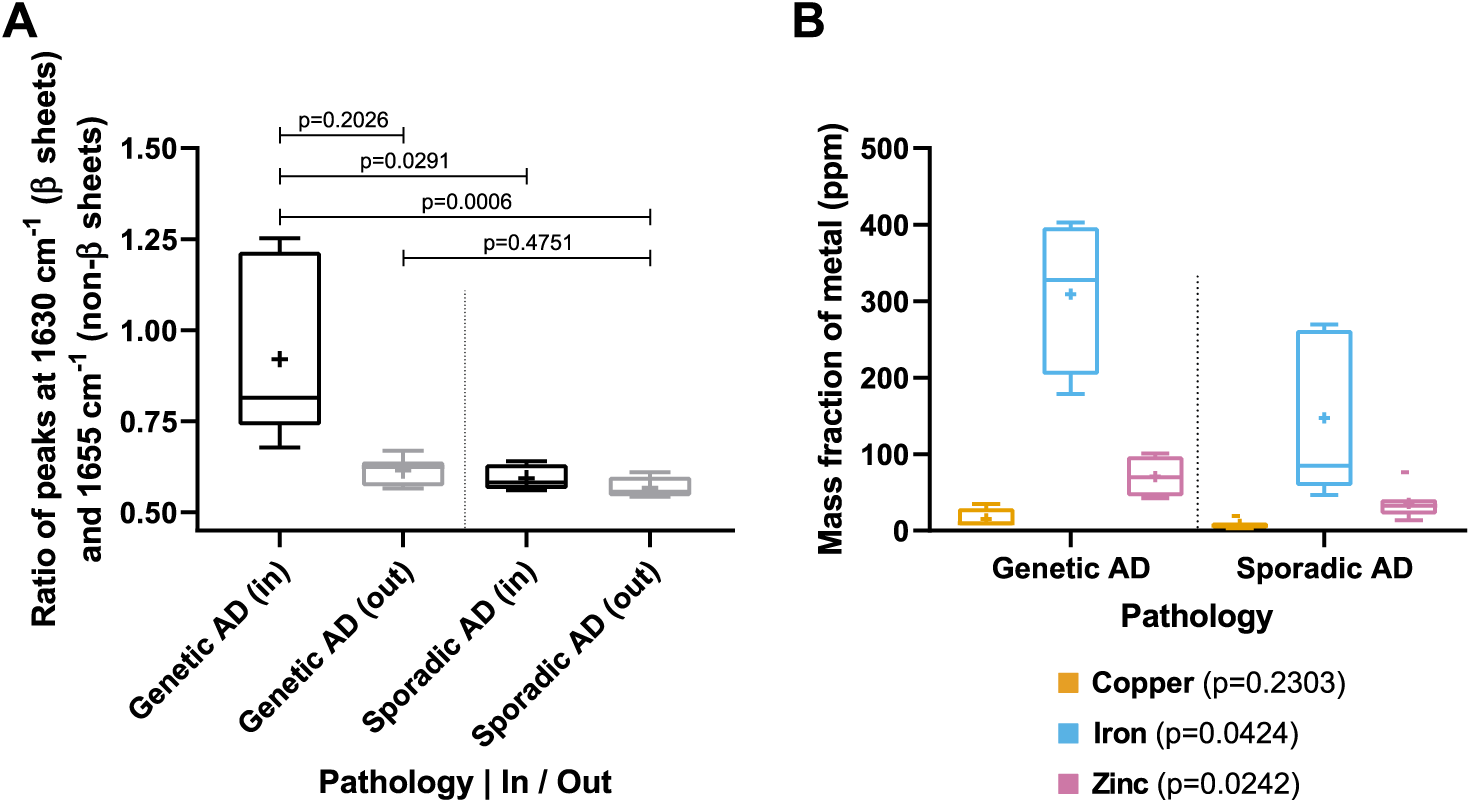
(A) Amyloid β-sheet content and (B) metal levels within Aβ plaques from Alzheimer’s Disease patients. β-sheet content is computed from peak ratios on µFTIR spectra and averaged per plaque (*n* = 6 for each patient). Metal levels are obtained by peak fitting on µXRF hyperspectra and averaged within a ROI on a plaque (*n* = 4 for the genetic AD, *n* = 7 for the sporadic AD).

Levels of metals within Aβ plaques (Figure 4, panel B) were higher (iron: +109 %; zinc: +97 %; copper: +88 %) for the genetic AD patient compared to the sporadic AD patient.

### 3.3. Transgenic AD models exhibit a variety of Aβ contrasts in XPCT

To further investigate the relationship between XPCT Aβ contrast and Aβ composition, we used 4 different transgenic rodent strains (Figure 5A): the J20 mice and the TgF344 rats had small hyperintense plaques (typically in the range 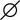25–30 µm for J20; and 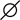15–20 µm for TgF344) mostly seen in the hippocampus (movies), while the APPPS1 and ArcAβ strains had numerous and large hypointense plaques (APPPS1: 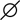40–45 µm; ArcAβ: 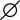45–60 µm), which however frequently exhibited a hyperintense core (APPPS1: 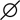10–15 µm; ArcAβ: 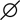15–20 µm), and were mainly located in the cortex but also in the hippocampus. Signal intensity measurements from ∼ 20 segmented plaques per sample are shown in Figure 5 B.

**Figure 5:**
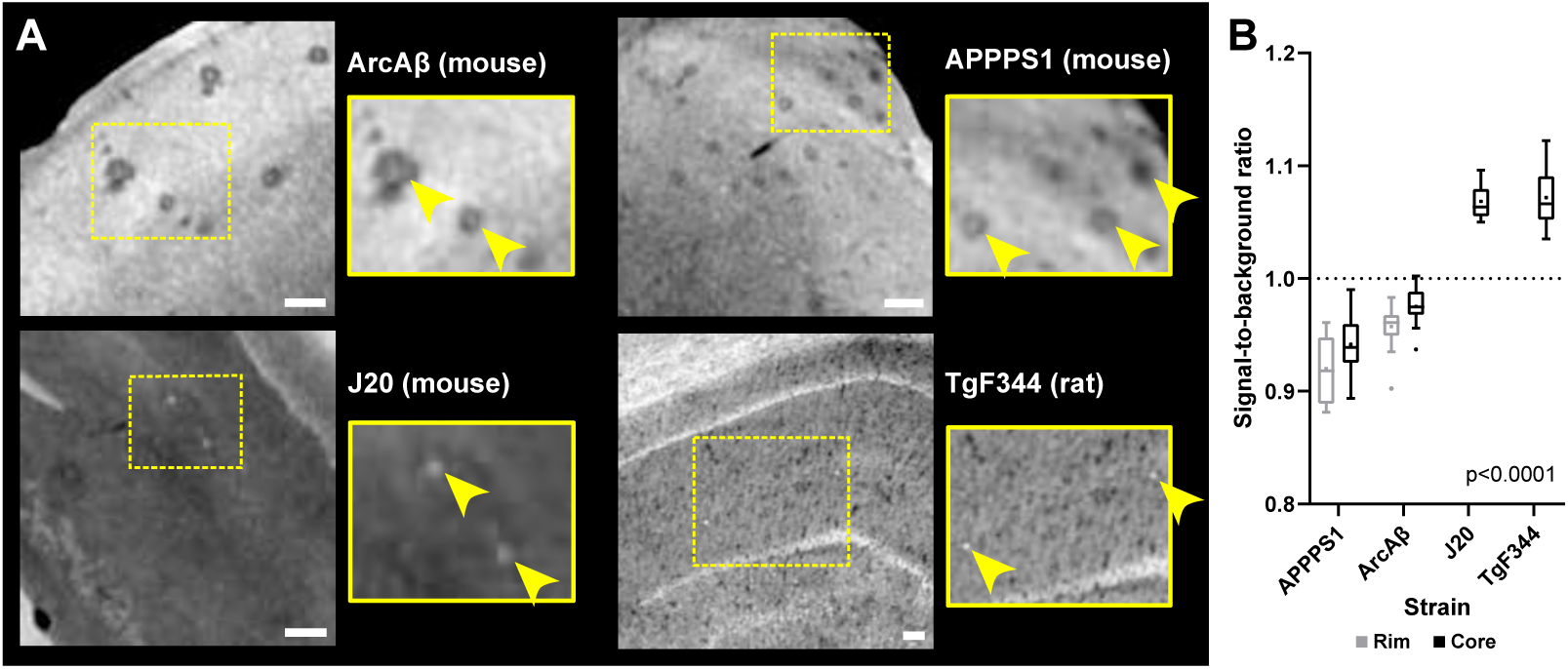
(A) XPCT slices of brain tissue with Aβ plaques from transgenic rodents and (B) associated contrasts of Aβ plaques. Insets are zoomed single-slice ROIs of the main views. Plaques from ArcAβ and APPPS1 mice are hypointense and mainly observed in the cortex. Plaques from J20 mice and TgF344 rats are hyperintense and mostly located in the hippocampus. Comparisons of signal-to-background ratio between hyperintense-plaque strains and hypointense-plaque strains yielded significant differences (*p* < 0.0001). Scale bars equal 100 µm.

### 3.4. β-sheet content is constant across transgenic strains while metal entrapment is highly variable

The contralateral hemispheres were used to study fibrillar and elemental composition of these plaques. The β-sheet content (Figure 6, A) increased in fluorescently-labelled spots in the rodent brain slices (measurements “In” vs. “Out” of fluorescent Aβ plaques: +37 % for APPPS1, +36 % for ArcAβ, +38 % for J20, +47 % for TgF344). The β-sheet content in the surrounding tissue (“Out”) in transgenic animals was similar to the level in corresponding wild-type animals (APPPS1: +6 %; ArcAβ: +2 %; J20: +9 %; TgF344: −6 %).

**Figure 6:**
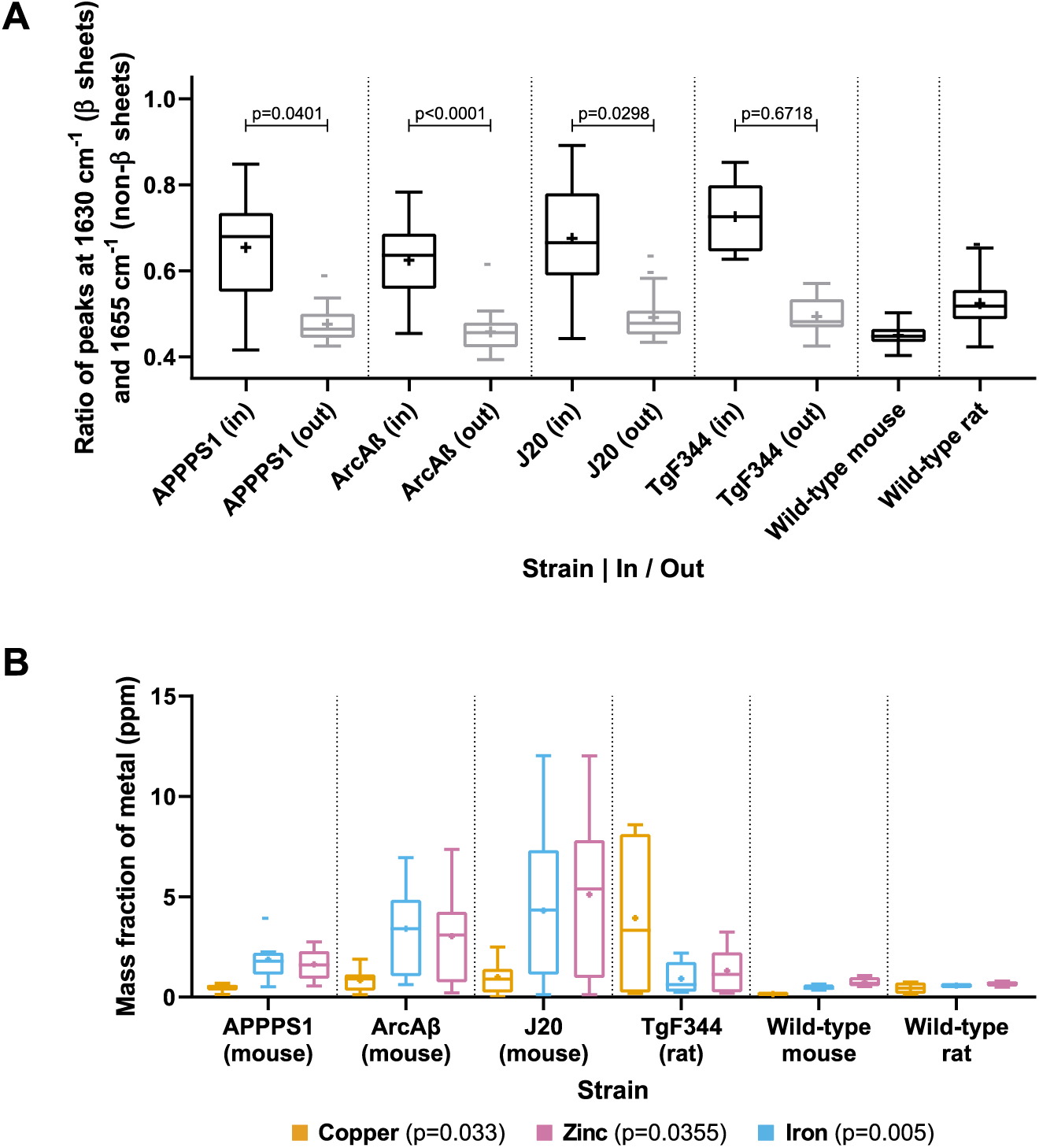
(A) Amyloid β-sheet content and (B) metal levels within Aβ plaques and in the surrounding tissue from transgenic rodents. β-sheet content is computed from peak ratios on µFTIR spectra and averaged per plaque (*n* = 9 − 20 plaques per strain). Metal levels are obtained by peak fitting on µXRF hyperspectra and averaged within a ROI on a plaque (n=4-17 plaques per strain).

All transgenic strains also had increased levels of iron, zinc and copper at the analyzed Aβ plaques compared to their wild-type counterparts. However, µXRF-measured metal composition of the Aβ plaques (Figure 6, B) varied between the different strains. J20 mice had the highest levels of zinc and iron compared to the other strains (zinc: +583 % vs. WT; iron: +803 % vs. WT) and TgF344 rats had the highest level of copper (+770 % vs. WT). Meanwhile, APPPS1 mice had the lowest mean levels and ArcAβ mice had intermediate levels.

### 3.5. Comparison between human and rodent datasets

Figure 7 shows a qualitative comparison of the multimodal dataset obtained with XPCT, µFTIR and µXRF, between transgenic rodents and humans. XPCT (left row) highlighted 3 distinct types of Aβ plaques: small hyperintense plaques in J20 mice and TgF344; larger, hypointense plaques with a bright core in ArcAβ and APPPS1; and similarly large, but diffusely hyperintense plaques in the genetic case of AD. All types of plaques exhibited an increased level of β-sheet content which matched the fluorescence signal, as measured by µFTIR (central row), and which tended to be increased in the human case compared to animal models. The accumulation of endogenous metals (right row) was mainly restricted to the (hyperintense) core of the plaque in rodents, while it appeared consistently diffuse over the whole µXRF FOV in the human plaques.

**Figure 7:**
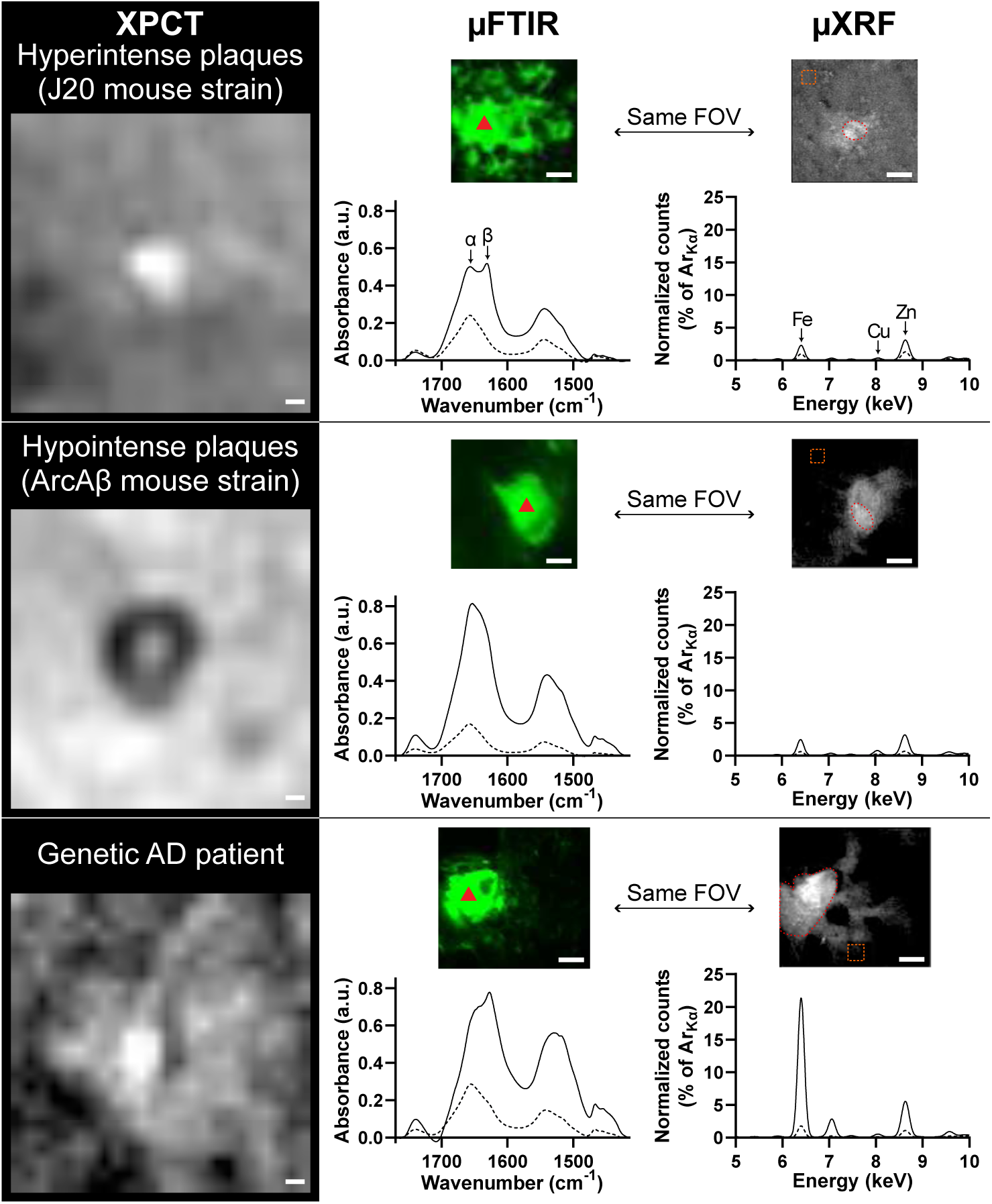
Multimodal dataset obtained with the three techniques on Aβ plaques. XPCT scans (left column) were acquired on an unsectioned sample, while µFTIR (center) & µXRF (right) hyperspectra were acquired on a paired, sectioned, sample. Inline brightfield microscopy was used to match the fields of view (FOVs) between µFTIR and µXRF. Representative measurements in the plaques are indicated with a red arrowhead (µFTIR) and red outline (µXRF), and plotted as plain lines. Measurements outside the plaques (µFTIR: acquired in fluorescence-free regions outside the shown FOVs; µXRF: orange outline) are plotted as dotted lines. In the case of µFTIR, a linear baseline correction was applied on the Amide I band, from 1415 cm^−1^ to 1770 cm^−1^. In the case of µXRF, maps show the summed µXRF signal and spectra were normalized with the K*_α_* peak of argon (Ar). Scale bars equal 10 µm.

## 4. Discussion

The present study sought to provide a rationale for the detection for Aβ plaques with synchrotron X-ray phase-contrast tomography, through a correlative approach between the local contrast — or signal intensity — and the fibrillar (µFTIR) and elemental (µXRF) compositions. The experimental design relied on three imaging modalities, all available at synchrotron SOLEIL, and included brain samples from patients diagnosed with Alzheimer’s Disease (AD) and transgenic rodents which developed amyloidosis. 2D correlative examination of Aβ plaques through µFTIR and µXRF has been an active field of research for 15 years (e.g. [15, 19, 35, 36]), which constantly highlighted that local elevations of β-sheet content and metal trapping were concomitant features across a variety of AD specimens, including in cellular models [37]. However this is the first time that the composition of Aβ plaques is linked with a 3D imaging method such as XPCT.

Using propagation-based imaging, now the most widely employed XPCT method for virtual histology of label-free samples, we made original observations. In animals, Aβ plaques from the J20 mouse strain and the TgF344 rat strain appeared as small bright (hyperintense) spots, as already reported for these and for other transgenic models such as 3xTg [7], APP/PS1 [6, 7], TauPS2APP [4] and B6C3-Tg [10]. But surprisingly, Aβ plaques in the newly imaged APPPS1 and ArcAβ mouse strains appeared as dark (hypointense) spots, with the largest plaques frequently exhibiting an hyperintense core. In humans, we confirmed previous observations that parenchymal Aβ plaques are usually not detected with XPCT, except rare deposits which were reported as calcified [13, 38]. However, we unveiled that Aβ contrast in the human samples is more pronounced in an inherited case of AD — of which the first XPCT images are reported here — compared to sporadic AD. This variable appearance of Aβ aggregates is reminiscent of Aβ polymorphism, which describes, both in animal models and in humans, the existence of subtypes of Aβ plaques [39, 40].

FTIR microspectroscopy can be used to quantify the β-sheet content in individual Aβ plaques with exquisite sensitivity. Indeed, the ratio of the β-sheet over non-β-sheet frequencies is used as a surrogate for fibrillar content. Aβ plaques were localized with a pan-amyloid fluorescent stain, and we found a relatively similar level of fibrils in all transgenic models (interquartile ranges of ratio were: 0.55–0.73 for APPPS1; 0.56–0.69 for ArcAβ; 0.59–0.78 for J20; 0.65–0.80 for TgF344). Not surprinsingly, the genetic AD patient had much higher levels of fibrils (interquartile ranges of ratio: 0.74–1.22) than the sporadic patient (0.54–0.60), probably due to the presence of more diffuse deposits in the sporadic case. Overall, these µFTIR results showed that the fibrillar content of Aβ plaques is not the sole factor responsible for XPCT detection.

Multiple studies — based on X-ray fluorescence [20] or other methods like inductively coupled plasma mass spectrometry (ICP-MS) [41, 42] — have linked AD pathogenesis with a disruption of the metal homeostasis in the brain, and more specifically copper, iron and zinc. These endogenous metals, whether in ionic or non-ionic forms [18], are locally trapped in Aβ plaques at the expense of the surrounding tissue [15]. The metal concentrations we detected in the human Aβ plaques matched those from the seminal study by Lovell et al. [43]. As already observed, animal models had much lower metal quantities associated with Aβ plaques than humans [19, 44]. Importantly, copper, zinc and iron levels in the genetic patient were all significantly higher (2-fold increase) compared to the sporadic AD patient. In animals, Aβ plaques from each strain preferentially bound to different metals (elevated Zn and Fe in J20; elevated Cu in TgF344). It has already been reported that iron is dysregulated within the hippocampus of J20 mouse model of AD [45]. And some differences between mouse and rat hippocampus — particularly on copper accumulation [46] — may explain our results on copper-enriched plaques in the rat model compared to mouse strain.

Overall, the combination of XPCT, µFTIR and µXRF (perfomed on the same brains with advanced amyloid pathology) allows us to propose the following interpretations. In mouse models, Aβ plaques form numerous well-defined, and clearly visible deposits on XPCT images. Elevated levels of metals, mainly restricted to a small core area on XRF maps, are associated with a positive signal (hyperintensity) detectable in all three strains on XPCT. Importantly, the brighter signals are detected in the strain with the highest level of metals (J20). However, a larger dark rim (hypointensity) extending this core is also detected in APPPS1 and ArcAβ, while absent in J20. Aβ fibrils, which match the fluorescence area and exceed the core area, cannot explain this difference as similar levels of fibrils were measured with µFTIR. Other factors, such as structural [40] or proteomic [47] differences between strains, might play a role.

In the transgenic rat, the pathology develops more slowly. Few plaques, representing only a subset of the global amyloid burden, are indeed detected with XPCT, as small hyperintense spots. In contrast, amyloid fluorescence was widely detected in hippocampus, which pointed to a spatially widespread pathology (Supplemental Figure 1). In line with this, additional µFTIR mapping experiments performed at ALBA synchrotron (beamline MIRAS), showed a global and unimodal increase in fibrillar content (Supplemental Figure 2), suggesting a diffuse accumulation of fibrils in the entire hippocampus. This implies that the few, large fluorescent areas that were targeted in our µFTIR/µXRF experiments, are likely to be biased and maybe not be entirely representative of the pathology. Nevertheless, if we assume that the largest fluorescent plaques are the ones that give rise to XPCT detection, the high amounts of copper highlighted by µXRF specifically in this model, are consistent with the results from mouse models, and could explain the bright appearance, restricted to the core of Aβ plaques in XPCT.

In humans, our data from two patients are obviously insufficient to disentangle the combined effect of β-sheet level and metal entrapment, which both drastically increased in the inherited case vs. sporadic one. However, the strikingly diffuse pattern of metal entrapment observed in XRF (in comparison to rodent models, Figure 7) seems consistent with the detection of hyperintense plaques on XPCT images. More samples would be needed to understand the local contrast in different types of Aβ plaques (diffuse, neuritic) which could not be properly discriminated here. A direct comparison, at the individual plaque level, between 3D XPCT and subsequent 2D µFTIR/µXRF would be ideal for that purpose, but would require cryogenic XPCT to avoid the chemical fixation used here (incompatible with µXRF). Also, tau pathology, which is present in human samples in the form of neurofibrillary tangles (and potentially associated with microcalcifications [48]), but lacking in the rodent strains studied here, could contribute to the XPCT contrast, as it was recently shown in a triple-transgenic mouse model [49]. It is nevertheless remarkable that Aβ plaques from the genetic case display a detectable XPCT contrast, and this observation deserves further confirmation in other familial cases of AD.

In conclusion, XPCT has attracted increasing interest in neuroscience these last years for its ability to provide valuable microstructural characterization (myelin, vessels, neuron density, protein deposits) in excised, intact, brain samples. XPCT can now be acquired in a few minutes at various resolutions, enabling a virtual dissection from the organ-level to the (sub-)cellular level. Indeed, high-resolution scans of whole organs are now possible as recently demonstrated by the particularly impressive 3D imaging of a full human brain [50]. The present report reinforces the potential of XPCT to capture key aspects of the Aβ pathology. However, if XPCT is envisioned as a potential screening method for Aβ pathology in brain samples, it is crucial to understand which factors drive the detection of some and not all Aβ plaques. This study was specifically designed to tackle this question, and highlighted sharp XPCT differences in Aβ pathology, which were related to the different levels of metals locally bound to Aβ plaques. This model-specific, dyshomeostasis of endogenous metals might be a downstream consequence of the genetic alterations, and could be region-specific or depend on the disease stage of progression. Additional factors, other than the overall β-sheet content, are likely to contribute to the XPCT contrast and remain to be identified.

## Acknowledgements

The authors acknowledge Synchrotron SOLEIL (Saint-Aubin, France) for allocation of beamtime within the scope of proposals #20200258 and #20220319 and we would like to thank the staff for assistance in using beamlines ANATOMIX (M. Scheel, J. Perrin), SMIS and NANOSCOPIUM (K. Medjoubi, A. Somogyi). ANATOMIX is an Equipment of Excellence (EQUIPEX) funded by the Investments for the Future program of the French National Research Agency (ANR), project NanoimagesX, grant no. ANR-11-EQPX-0031. We would like to thank Clément Tavakoli (STROBE, Univ. Grenoble-Alpes, Grenoble; Univ. Lyon 1, Lyon) for his help during the acquisitions at SOLEIL. The study was further supported by the ALBA Synchrotron Light Source (ALBA-CELLS, Barcelona, Spain; beamline MIRAS; proposal 2018082957). This study was performed within the framework of LABEX PRIMES (ANR-11-LABX-0063) of Université de Lyon, within the “Investissements d’Avenir” program (ANR-11-IDEX-0007) of the French National Research Agency (ANR). We are grateful to Corinne Perrin from the Tumorothèque Est tissue bank, CRB-HCL (Lyon, France), for managing human brain samples. We also thank Jérémy Verchère (ANSES, Lyon, France) for the management of APPPS1 brains. p-FTAA was a gift from the laboratory of chemistry at ENS-Lyon. S. Singh received support from the Auvergne-Rhône-Alpes region (FUI GigaQuant project).

## Authors’ contributions (according to Contributor Role Taxonomy (CRediT))

- Conceptualization: FC
- Methodology: EB, SB, FC
- Resources: CS, TW, TD, DM, TB (APPPS1), JK (ArcAβ), NR (J20), HB (TgF344), CO, EB, SB
- Investigation: MC, CS, TW, TD, CO, MW, EB, SB
- Formal analysis: MC, CS, SS, SB
- Data curation: MC, CS, SB
- Software:
- Visualization: MC, DM, FC
- Validation: MC
- Supervision: CS, EB, SB, FC
- Project administration: FC
- Funding acquisition: FC
- Writing – original draft: MC, FC
- Writing – review & editing: MC, DM, CS, TW, TD, JK, MW, EB, FC

## Supplemental Document — Why are amyloid-β plaques detected by X-ray phase-contrast imaging? Role of metals revealed through combined synchrotron infrared and X-ray fluorescence microscopies

**Supplemental Figure 1:**
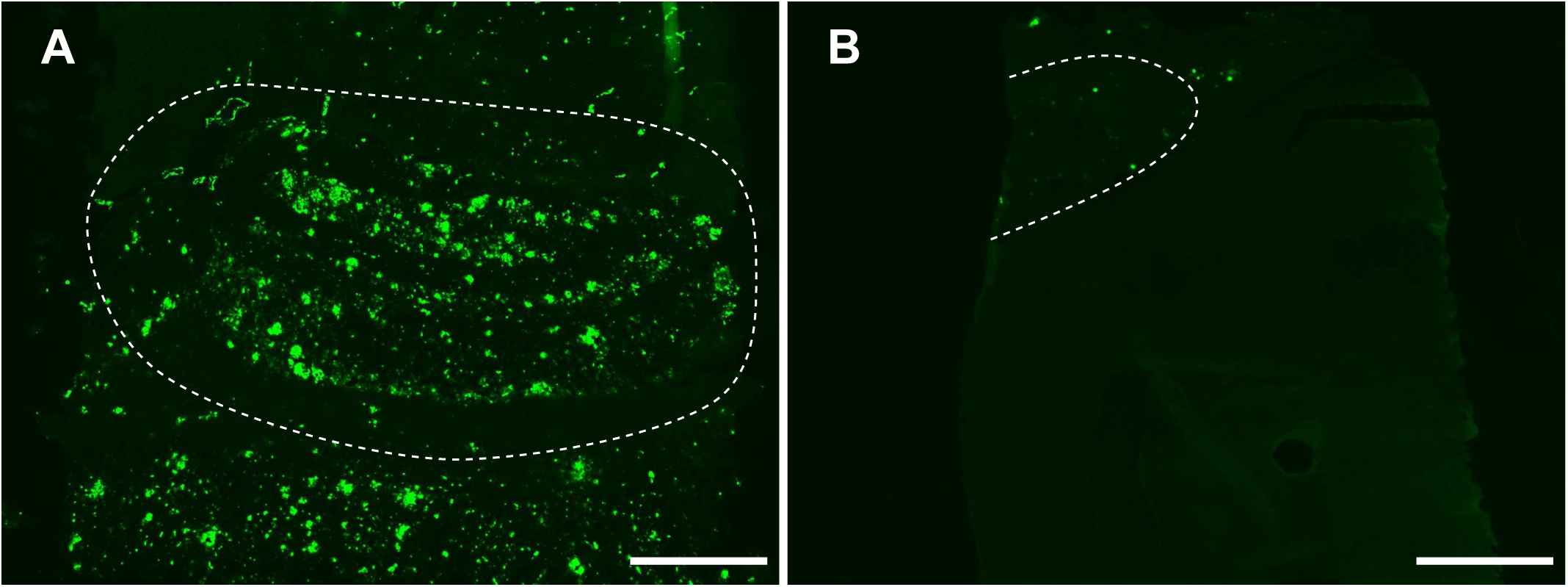
Fluorescence microscopy images of p-FTAA-stained sections of a TgF344 rat brain (A) and a J20 mouse brain (B), near hippocampus (shown with dotted line). Both sections were acquired with the same exposure time and rendered with the same value range. One can notice the p-FTAA staining reveals more diffuse structures in the rat brain. Scale bars equal 1000 µm.

### A Extra parameters for synchrotron experiments

**Supplemental Table 1:** Acquisition and reconstruction parameters for XPCT (ANATOMIX) per session

| Numerical parameter | Values: | Session #1 | Session #2 |
| --- | --- | --- | --- |
| Undulator gap (mm) |  | 11.7 | 9.7 |
| Exposure time per projection (ms) |  | 200 | 50 |
| Gold filter thickness ($\mu\text{m}$) | | 20 | 20 |
| Detector-to-sample distance (cm) |  | 113 | 30 |
| Magnification |  | 1 × | 2.1 × |
| Projection pixel size ($\mu\text{m}$) | | 6.5 | 3.09 |
| Number of projections |  | 2600 | 4000 |
| Weight of unsharp mask filter |  | 0.5 | 0.5 |
| Radius of unsharp mask filter (px) |  | 2 | 2 |
| $\delta/\beta$ for Paganin filtering | | 1525 | 1400 |

**Supplemental Table 2:** Acquisition parameters for µFTIR (SMIS)

| Parameter | Value |
| --- | --- |
| Mode | Dual aperture |
| Apodization | Strong Norton & Beer |
| Objective type | Schwarzschild |

Supplemental Table 2: Acquisition parameters for $\mu\text{FTIR}$ (SMIS)
| Numerical parameter | Value |
| --- | --- |
| Spectral resolution ($\text{cm}^{-1}$) | 4 |
| Zero-fill factor | 2 |
| Aperture size ($\mu\text{m}^2$) | $10 \times 10$ |
| Co-addition | 256 |
| Objective numerical aperture | 0.65 |
| Objective magnification | 32 |

**Supplemental Table 3:**
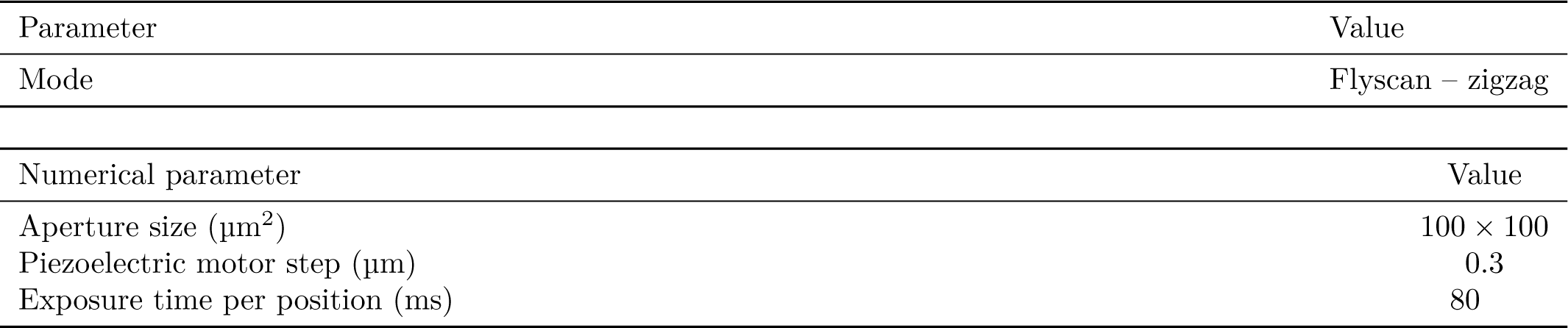
Acquisition parameters for µXRF (NANOSCOPIUM)

### B Scanning the same positions on SMIS and NANOSCOPIUM

Scanning the same positions on SMIS and NANOSCOPIUM was crucial to associate the FTIR and XRF measures. However, beamtimes were two weeks apart and required different sample holders and orientations. To overcome this challenge, two coordinate systems 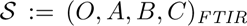 and 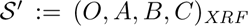 were created using the inner corners of the membrane, so that the new points 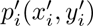 were obtained from *p_i_*(*x_i_, y_i_*) by solving

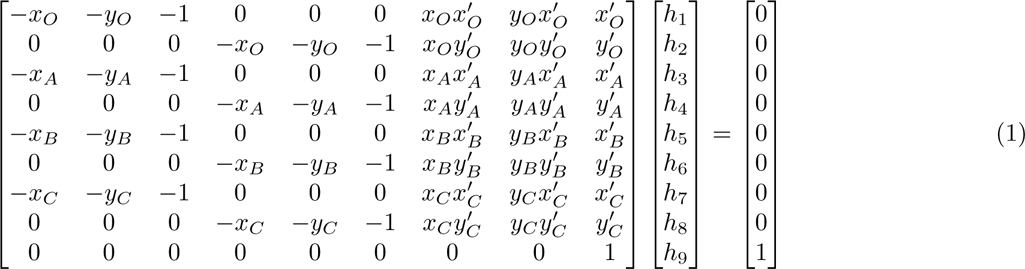

then

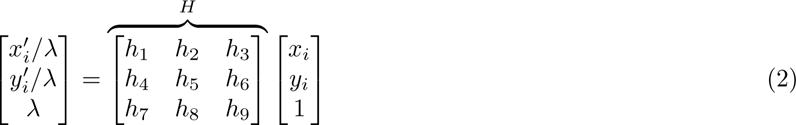

where *H* is the homography matrix and *λ* is a factor.

### C Measures at ALBA synchrotron (Barcelona, Spain)

Large-field FTIR images were obtained using a 64 × 64 px full-field focal plane array (FPA) detector cooled with liquid nitrogen. Images were taken by FPA detector on areas of 210 × 210 µm^2^ with 128 accumulated spectra per point at 8 cm^−1^ spectral resolution and 10 µm pixel size. The spectra were background-corrected using extracellular regions close to the tissue. From each IR image, the total absorbance cartogram was generated by integrating at specific peaks in amide I area (1600 cm^−1^–1700 cm^−1^), i.e. the ratio of peaks characteristic for β-sheet structures absorbing at 1625 cm^−1^ and α-helix structures assigned at 1655 cm^−1^ (Supplemental Figure 2).

**Supplemental Figure 2:**
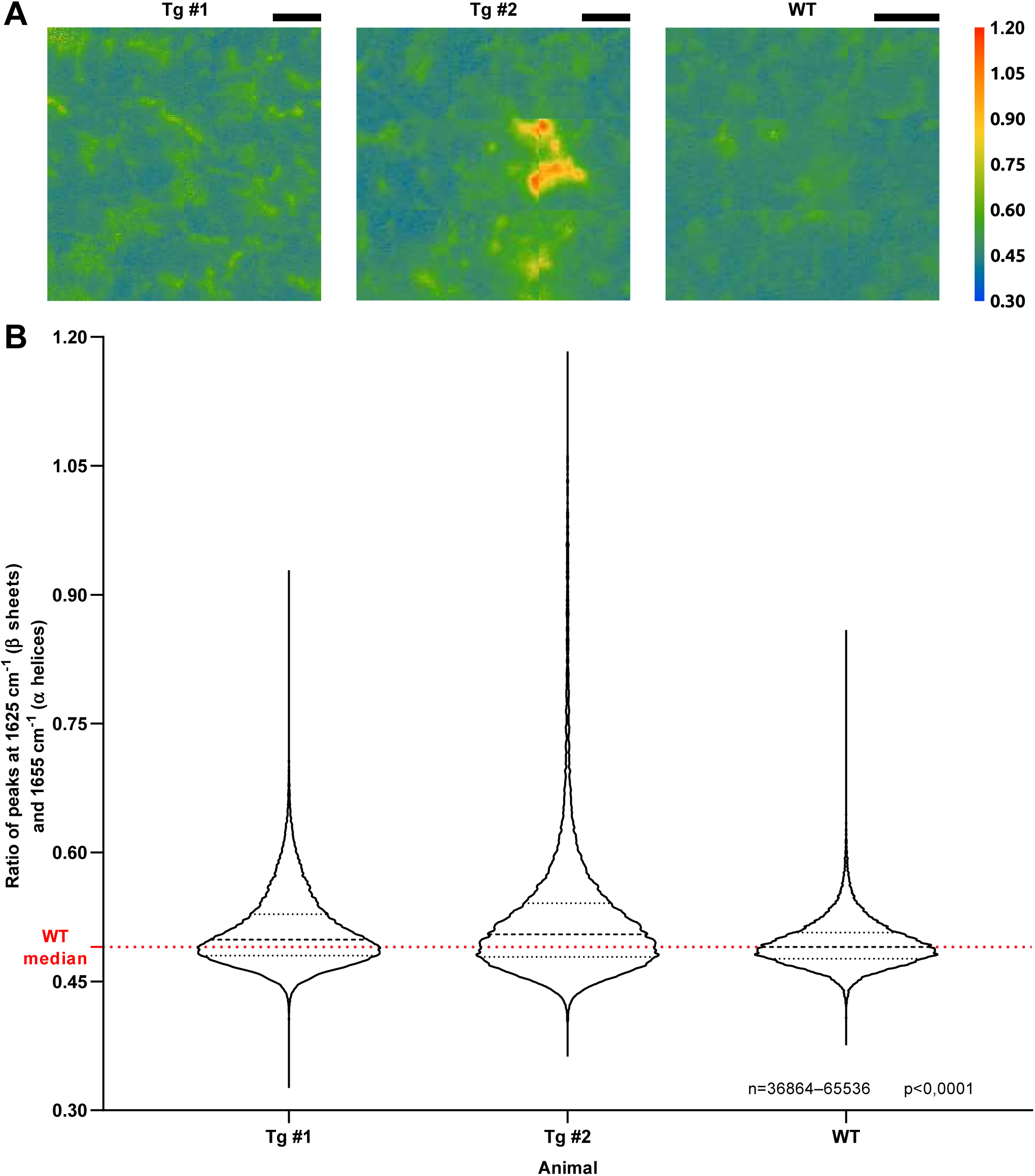
(A) FPA-FTIR maps of the β-sheet content within the hippocampus and (B) measures from TgF344 rats (Tg #1 and Tg #2) and a wild-type rat (WT), acquired at ALBA synchrotron. Mean and standard deviation of Tg #1: 0.508 ± 0.041; Tg #2: 0.523 ± 0.078; WT: 0.494 ± 0.026. Scale bars equal 50 µm.

## References

[1] M. A. Kolanko, Z. Win, F. Loreto, N. Patel, C. Carswell, A. Gontsarova, R. J. Perry, P. A. Malhotra, Amyloid PET imaging in clinical practice, Practical Neurology 20 (2020) 451–462. doi:10.1136/practneurol-2019-002468.

[2] S. S. Mirra, A. Heyman, D. McKeel, S. M. Sumi, B. J. Crain, L. M. Brownlee, F. S. Vogel, J. P. Hughes, G. v. Belle, L. Berg, participating CERAD neuropathologists, The Consortium to Establish a Registry for Alzheimer’s Disease (CERAD): Part II. Standardization of the neuropathologic assessment of Alzheimer’s disease, Neurology 41 (1991) 479–479. doi:10.1212/WNL.41.4.479.

[3] D. R. Thal, U. Rüb, M. Orantes, H. Braak, Phases of A beta-deposition in the human brain and its relevance for the development of AD, Neurology 58 (2002) 1791–1800. doi:10.1212/wnl.58.12.1791.

[4] A. Astolfo, A. Lathuilière, V. Laversenne, B. Schneider, M. Stampanoni, Amyloid-*β* plaque deposition measured using propagation-based X-ray phase contrast CT imaging, Journal of Synchrotron Radiation 23 (2016) 813–819. doi:10.1107/S1600577516004045.

[5] L. Massimi, I. Bukreeva, G. Santamaria, M. Fratini, A. Corbelli, F. Brun, S. Fumagalli, L. Maugeri, A. Pacureanu, P. Cloetens, N. Pieroni, F. Fiordaliso, G. Forloni, A. Uccelli, N. Kerlero de Rosbo, C. Balducci, A. Cedola, Exploring Alzheimer’s disease mouse brain through X-ray phase contrast tomography: From the cell to the organ, NeuroImage 184 (2019) 490–495. doi:10.1016/j.neuroimage.2018.09.044.

[6] L. Massimi, N. Pieroni, L. Maugeri, M. Fratini, F. Brun, I. Bukreeva, G. Santamaria, V. Medici, T. E. Poloni, C. Balducci, A. Cedola, Assessment of plaque morphology in Alzheimer’s mouse cerebellum using three-dimensional X-ray phase-based virtual histology, Scientific reports 10 (2020) 11233. doi:10.1038/s41598-020-68045-8.

[7] M. Chourrout, M. Roux, C. Boisvert, C. Gislard, D. Legland, I. Arganda-Carreras, C. Olivier, F. Peyrin, H. Boutin, N. Rama, T. Baron, D. Meyronet, E. Brun, H. Rositi, M. Wiart, F. Chauveau, Brain virtual histology with X-ray phase-contrast tomography Part II: 3D morphologies of amyloid-*β* plaques in Alzheimer’s disease models, Biomedical Optics Express 13 (2022) 1640. doi:10.1364/BOE.438890.

[8] M. Töpperwien, M. Krenkel, D. Vincenz, F. Stöber, A. M. Oelschlegel, J. Goldschmidt, T. Salditt, Three-dimensional mouse brain cytoarchitecture revealed by laboratory-based x-ray phase-contrast tomography, Scientific Reports 7 (2017) 42847. doi:10.1038/srep42847.

[9] K. Noda-Saita, A. Yoneyama, Y. Shitaka, Y. Hirai, K. Terai, J. Wu, T. Takeda, K. Hyodo, N. Osakabe, T. Yamaguchi, M. Okada, Quantitative analysis of amyloid plaques in a mouse model of Alzheimer’s disease by phase-contrast X-ray computed tomography, Neuroscience 138 (2006) 1205–1213. doi:10.1016/j.neuroscience.2005.12.036.

[10] D. M. Connor, H. Benveniste, F. A. Dilmanian, M. F. Kritzer, L. M. Miller, Z. Zhong, Computed tomography of amyloid plaques in a mouse model of Alzheimer’s disease using diffraction enhanced imaging, NeuroImage 46 (2009) 908–914. doi:10.1016/j.neuroimage.2009.03.019.

[11] B. Pinzer, M. Cacquevel, P. Modregger, S. McDonald, J. Bensadoun, T. Thuering, P. Aebischer, M. Stampanoni, Imaging brain amyloid deposition using grating-based differential phase contrast tomography, NeuroImage 61 (2012) 1336–1346. doi:10.1016/j.neuroimage.2012.03.029.

[12] J. Albers, S. Pacilé, M. A. Markus, M. Wiart, G. Vande Velde, G. Tromba, C. Dullin, X-ray-Based 3D Virtual Histology-Adding the Next Dimension to Histological Analysis, Molecular Imaging and Biology 20 (2018) 732–741. doi:10.1007/s11307-018-1246-3.

[13] M. Eckermann, B. Schmitzer, F. van der Meer, J. Franz, O. Hansen, C. Stadelmann, T. Salditt, Three-dimensional virtual histology of the human hippocampus based on phase-contrast computed tomography, Proceedings of the National Academy of Sciences 118 (2021) e2113835118. doi:10.1073/pnas.2113835118.

[14] M. Chourrout, H. Rositi, E. Ong, V. Hubert, A. Paccalet, L. Foucault, A. Autret, B. Fayard, C. Olivier, R. Bolbos, F. Peyrin, C. Crola-da-Silva, D. Meyronet, O. Raineteau, H. Elleaume, E. Brun, F. Chauveau, M. Wiart, Brain virtual histology with X-ray phase-contrast tomography Part I: Whole-brain myelin mapping in white-matter injury models, Biomedical Optics Express 13 (2022) 1620. doi:10.1364/BOE.438832.

[15] L. M. Miller, Q. Wang, T. P. Telivala, R. J. Smith, A. Lanzirotti, J. Miklossy, Synchrotron-based infrared and X-ray imaging shows focalized accumulation of Cu and Zn co-localized with *β*-amyloid deposits in Alzheimer’s disease, Journal of Structural Biology 155 (2006) 30–37. doi:10.1016/j.jsb.2005.09.004.

[16] M. Rak, M. R. D. Bigio, S. Mai, D. Westaway, K. Gough, Dense-core and diffuse A*β* plaques in TgCRND8 mice studied with synchrotron FTIR microspectroscopy, Biopolymers 87 (2007) 207–217. doi:10.1002/bip.20820.

[17] C. R. Liao, M. Rak, J. Lund, M. Unger, E. Platt, B. C. Albensi, C. J. Hirschmugl, K. M. Gough, Synchrotron FTIR reveals lipid around and within amyloid plaques in transgenic mice and Alzheimer’s disease brain, The Analyst 138 (2013) 3991. doi:10.1039/c3an00295k.

[18] J. Everett, F. Lermyte, J. Brooks, V. Tjendana-Tjhin, G. Plascencia-Villa, I. Hands-Portman, J. M. Donnelly, K. Billimoria, G. Perry, X. Zhu, P. J. Sadler, P. B. O’Connor, J. F. Collingwood, N. D. Telling, Biogenic metallic elements in the human brain?, Science Advances 7 (2021) eabf6707. doi:10.1126/sciadv.abf6707.

[19] A. C. Leskovjan, A. Lanzirotti, L. M. Miller, Amyloid plaques in PSAPP mice bind less metal than plaques in human Alzheimer’s disease, NeuroImage 47 (2009) 1215–1220. doi:10.1016/j.neuroimage.2009.05.063.

[20] S. A. James, Q. I. Churches, M. D. de Jonge, I. E. Birchall, V. Streltsov, G. McColl, P. A. Adlard, D. J. Hare, Iron, Copper, and Zinc Concentration in A*β* Plaques in the APP/PS1 Mouse Model of Alzheimer’s Disease Correlates with Metal Levels in the Surrounding Neuropil, ACS Chemical Neuroscience 8 (2017) 629–637. doi:10.1021/acschemneuro.6b00362.

[21] A. C. Leskovjan, A. Kretlow, A. Lanzirotti, R. Barrea, S. Vogt, L. M. Miller, Increased brain iron coincides with early plaque formation in a mouse model of Alzheimer’s disease, NeuroImage 55 (2011) 32–38. doi:10.1016/j.neuroimage.2010.11.073.

[22] I. Alafuzoff, D. R. Thal, T. Arzberger, N. Bogdanovic, S. Al-Sarraj, I. Bodi, S. Boluda, O. Bugiani, C. Duyckaerts, E. Gelpi, S. Gentleman, G. Giaccone, M. Graeber, T. Hortobagyi, R. Höftberger, P. Ince, J. W. Ironside, N. Kavantzas, A. King, P. Korkolopoulou, G. G. Kovács, D. Meyronet, C. Monoranu, T. Nilsson, P. Parchi, E. Patsouris, M. Pikkarainen, T. Revesz, A. Rozemuller, D. Seilhean, W. Schulz-Schaeffer, N. Streichenberger, S. B. Wharton, H. Kretzschmar, Assessment of *β*-amyloid deposits in human brain: A study of the Brain-Net Europe Consortium, Acta Neuropathologica 117 (2009) 309–320. doi:10.1007/s00401-009-0485-4.

[23] M. Knobloch, U. Konietzko, D. C. Krebs, R. M. Nitsch, Intracellular A*β* and cognitive deficits precede *β*-amyloid deposition in transgenic arcA*β* mice, Neurobiology of Aging 28 (2007) 1297–1306. doi:10.1016/j.neurobiolaging.2006.06.019.

[24] R. Radde, T. Bolmont, S. A. Kaeser, J. Coomaraswamy, D. Lindau, L. Stoltze, M. E. Calhoun, F. Jäggi, H. Wolburg, S. Gengler, C. Haass, B. Ghetti, C. Czech, C. Hölscher, P. M. Mathews, M. Jucker, A*β*42-driven cerebral amyloidosis in transgenic mice reveals early and robust pathology, EMBO reports 7 (2006) 940–946. doi:10.1038/sj.embor.7400784.

[25] L. Mucke, E. Masliah, G.-Q. Yu, M. Mallory, E. M. Rockenstein, G. Tatsuno, K. Hu, D. Kholodenko, K. Johnson-Wood, L. McConlogue, High-Level Neuronal Expression of A*β*1-42 in Wild-Type Human Amyloid Protein Precursor Transgenic Mice: Synaptotoxicity without Plaque Formation, The Journal of Neuroscience 20 (2000) 4050–4058. doi:10.1523/JNEUROSCI.20-11-04050.2000.

[26] R. M. Cohen, K. Rezai-Zadeh, T. M. Weitz, A. Rentsendorj, D. Gate, I. Spivak, Y. Bholat, V. Vasilevko, C. G. Glabe, J. J. Breunig, P. Rakic, H. Davtyan, M. G. Agadjanyan, V. Kepe, J. R. Barrio, S. Bannykh, C. A. Szekely, R. N. Pechnick, T. Town, A Transgenic Alzheimer Rat with Plaques, Tau Pathology, Behavioral Impairment, Oligomeric A*β*, and Frank Neuronal Loss, Journal of Neuroscience 33 (2013) 6245–6256. doi:10.1523/JNEUROSCI.3672-12.2013.

[27] A. Åslund, C. J. Sigurdson, T. Klingstedt, S. Grathwohl, T. Bolmont, D. L. Dickstein, E. Glimsdal, S. Prokop, M. Lindgren, P. Konradsson, D. M. Holtzman, P. R. Hof, F. L. Heppner, S. Gandy, M. Jucker, A. Aguzzi, P. Hammarström, K. P. R. Nilsson, Novel Pentameric Thiophene Derivatives for in Vitro and in Vivo Optical Imaging of a Plethora of Protein Aggregates in Cerebral Amyloidoses, ACS Chemical Biology 4 (2009) 673–684. doi:10.1021/cb900112v.

[28] T. Weitkamp, M. Scheel, J. Perrin, G. Daniel, A. King, L. Roux, V, J.-L. Giorgetta, A. Carcy, F. Langlois, K. Desjardins, C. Menneglier, M. Cerato, C. Engblom, G. Cauchon, T. Moreno, C. Rivard, Y. Gohon, F. Polack, Microtomography on the ANATOMIX beamline at Synchrotron SOLEIL, 2022. arXiv:2002.03242.

[29] A. Mirone, E. Brun, E. Gouillart, P. Tafforeau, J. Kieffer, The PyHST2 hybrid distributed code for high speed tomographic reconstruction with iterative reconstruction and a priori knowledge capabilities, Nuclear Instruments and Methods in Physics Research Section B: Beam Interactions with Materials and Atoms 324 (2014) 41–48. doi:10.1016/j.nimb.2013.09.030.

[30] D. Legland, I. Arganda-Carreras, P. Andrey, MorphoLibJ: Integrated library and plugins for mathematical morphology with ImageJ, Bioinformatics (2016) btw413. doi:10.1093/bioinformatics/btw413.

[31] M. Toplak, S. T. Read, C. Sandt, F. Borondics, Quasar: Easy Machine Learning for Biospectroscopy, Cells 10 (2021) 2300. doi:10.3390/cells10092300.

[32] P. Bassan, H. J. Byrne, F. Bonnier, J. Lee, P. Dumas, P. Gardner, Resonant Mie scattering in infrared spectroscopy of biological materials – understanding the ‘dispersion artefact’, The Analyst 134 (2009) 1586. doi:10.1039/b904808a.

[33] D. Röhr, B. D. C. Boon, M. Schuler, K. Kremer, J. J. M. Hoozemans, F. H. Bouwman, S. F. El-Mashtoly, A. Nabers, F. Großerueschkamp, A. J. M. Rozemuller, K. Gerwert, Label-free vibrational imaging of different A*β* plaque types in Alzheimer’s disease reveals sequential events in plaque development, Acta Neuropathologica Communications 8 (2020) 222. doi:10.1186/s40478-020-01091-5.

[34] V. Solé, E. Papillon, M. Cotte, P. Walter, J. Susini, A multiplatform code for the analysis of energy-dispersive X-ray fluorescence spectra, Spectrochimica Acta Part B: Atomic Spectroscopy 62 (2007) 63–68. doi:10.1016/j.sab.2006.12.002.

[35] K. L. Summers, N. Fimognari, A. Hollings, M. Kiernan, V. Lam, R. J. Tidy, D. Paterson, M. J. Tobin, R. Takechi, G. N. George, I. J. Pickering, J. C. Mamo, H. H. Harris, M. J. Hackett, A Multimodal Spectroscopic Imaging Method To Characterize the Metal and Macromolecular Content of Proteinaceous Aggregates (“Amyloid Plaques”), Biochemistry 56 (2017) 4107–4116. doi:10.1021/acs.biochem.7b00262.

[36] A. D. Surowka, M. Czyzycki, A. Ziomber-Lisiak, A. Migliori, M. Szczerbowska-Boruchowska, On 2D-FTIR-XRF microscopy – A step forward correlative tissue studies by infrared and hard X-ray radiation, Ultramicroscopy 232 (2022) 113408. doi:10.1016/j.ultramic.2021.113408.

[37] N. Gustavsson, A. Paulus, I. Martinsson, A. Engdahl, K. Medjoubi, K. Klementiev, A. Somogyi, T. Deierborg, F. Borondics, G. K. Gouras, O. Klementieva, Correlative optical photothermal infrared and X-ray fluorescence for chemical imaging of trace elements and relevant molecular structures directly in neurons, Light: Science & Applications 10 (2021) 151. doi:10.1038/s41377-021-00590-x.

[38] M. Töpperwien, F. van der Meer, C. Stadelmann, T. Salditt, Correlative x-ray phase-contrast tomography and histology of human brain tissue affected by Alzheimer’s disease, NeuroImage 210 (2020) 116523. doi:10.1016/j.neuroimage.2020.116523.

[39] G. Di Fede, M. Catania, E. Maderna, R. Ghidoni, L. Benussi, E. Tonoli, G. Giaccone, F. Moda, A. Paterlini, I. Campagnani, S. Sorrentino, L. Colombo, A. Kubis, E. Bistaffa, B. Ghetti, F. Tagliavini, Molecular subtypes of Alzheimer’s disease, Scientific Reports 8 (2018) 1–14. doi:10.1038/s41598-018-21641-1.

[40] J. Rasmussen, J. Mahler, N. Beschorner, S. A. Kaeser, L. M. Häsler, F. Baumann, S. Nyström, E. Portelius, K. Blennow, T. Lashley, N. C. Fox, D. Sepulveda-Falla, M. Glatzel, A. L. Oblak, B. Ghetti, K. P. R. Nilsson, P. Hammarström, M. Staufenbiel, L. C. Walker, M. Jucker, Amyloid polymorphisms constitute distinct clouds of conformational variants in different etiological subtypes of Alzheimer’s disease, Proceedings of the National Academy of Sciences 114 (2017) 13018–13023. doi:10.1073/pnas.1713215114.

[41] D. J. Hare, J. K. Lee, A. D. Beavis, A. van Gramberg, J. George, P. A. Adlard, D. I. Finkelstein, P. A. Doble, Three-Dimensional Atlas of Iron, Copper, and Zinc in the Mouse Cerebrum and Brainstem, Analytical Chemistry 84 (2012) 3990–3997. doi:10.1021/ac300374x.

[42] M. Cruz-Alonso, B. Fernandez, A. Navarro, S. Junceda, A. Astudillo, R. Pereiro, Laser ablation ICP-MS for simultaneous quantitative imaging of iron and ferroportin in hippocampus of human brain tissues with Alzheimer’s disease, Talanta 197 (2019) 413–421. doi:10.1016/j.talanta.2019.01.056.

[43] M. Lovell, J. Robertson, W. Teesdale, J. Campbell, W. Markesbery, Copper, iron and zinc in Alzheimer’s disease senile plaques, Journal of the Neurological Sciences 158 (1998) 47–52. doi:10.1016/S0022-510X(98)00092-6.

[44] X. Zhu, T. W. Victor, A. Ambi, J. K. Sullivan, J. Hatfield, F. Xu, L. M. Miller, W. E. Van Nostrand, Copper accumulation and the effect of chelation treatment on cerebral amyloid angiopathy compared to parenchymal amyloid plaques, Metallomics 12 (2020) 539–546. doi:10.1039/c9mt00306a.

[45] D. W. Dekens, P. J. W. Naudé, J. N. Keijser, A. S. Boerema, P. P. De Deyn, U. L. M. Eisel, Lipocalin 2 contributes to brain iron dysregulation but does not affect cognition, plaque load, and glial activation in the J20 Alzheimer mouse model, Journal of Neuroinflammation 15 (2018) 330. doi:10.1186/s12974-018-1372-5.

[46] B. Sullivan, G. Robison, Y. Pushkar, J. K. Young, K. F. Manaye, Copper accumulation in rodent brain astrocytes: A species difference, Journal of Trace Elements in Medicine and Biology 39 (2017) 6–13. doi:10.1016/j.jtemb.2016.06.011.

[47] E. Drummond, T. Kavanagh, G. Pires, M. Marta-Ariza, E. Kanshin, S. Nayak, A. Faustin, V. Berdah, B. Ueberheide, T. Wisniewski, The amyloid plaque proteome in early onset Alzheimer’s disease and Down syndrome, Acta Neuropathologica Communications 10 (2022) 53. doi:10.1186/s40478-022-01356-1.

[48] E. Tsolaki, L. Csincsik, J. Xue, I. Lengyel, S. Bertazzo, Nuclear and cellular, micro and nano calcification in Alzheimer’s disease patients and correlation to phosphorylated Tau, Acta Biomaterialia 143 (2022) 138–144. doi:10.1016/j.actbio.2022.03.003.

[49] G. E. Barbone, A. Bravin, A. Mittone, A. Pacureanu, G. Mascio, P. Di Pietro, M. J. Kraiger, M. Eckermann, M. Romano, M. Hrabě de Angelis, P. Cloetens, V. Bruno, G. Battaglia, P. Coan, X-ray multiscale 3D neuroimaging to quantify cellular aging and neurodegeneration postmortem in a model of Alzheimer’s disease, European Journal of Nuclear Medicine and Molecular Imaging (2022). doi:10.1007/s00259-022-05896-5.

[50] C. L. Walsh, P. Tafforeau, W. L. Wagner, D. J. Jafree, A. Bellier, C. Werlein, M. P. Kühnel, E. Boller, S. Walker-Samuel, J. L. Robertus, D. A. Long, J. Jacob, S. Marussi, E. Brown, N. Holroyd, D. D. Jonigk, M. Ackermann, P. D. Lee, Imaging intact human organs with local resolution of cellular structures using hierarchical phase-contrast tomography, Nature Methods 18 (2021) 1532–1541. doi:10.1038/s41592-021-01317-x.

